# Intertwined Functions of Separase and Caspase in Cell Division and Programmed Cell Death

**DOI:** 10.1101/653584

**Authors:** Pan-Young Jeong, Ashish Kumar, Pradeep Joshi, Joel H. Rothman

**Affiliations:** Department of Molecular, Cellular, and Developmental Biology, and Neuroscience Research Institute, University of California, Santa Barbara, Santa Barbara, CA 93106, USA

**Keywords:** *C. elegans*, chromosome separation, separase, caspase, programmed cell death

## Abstract

Timely sister chromatid separation, promoted by separase, is essential for faithful chromosome segregation. Separase is a member of the CD clan of cysteine proteases, which also includes the pro-apoptotic enzymes known as caspases. We report that the *C. elegans* separase SEP-1, primarily known for its role in cell division, is required for apoptosis when the predominant pro-apoptotic caspase CED-3 is compromised. Loss of SEP-1 results in extra surviving cells in a weak *ced-3(−)* mutant, and suppresses the embryonic lethality of a mutant defective for the apoptotic suppressor *ced-9*/*Bcl-2*. We also report apparent non-apoptotic roles for CED-3 in promoting germ cell proliferation and germline meiotic chromosome disjunction and the normal rate of embryonic development. Moreover, loss of the soma-specific (CSP-3) and germline-specific (CSP-2) caspase inhibitors results in CED-3-dependent suppression of embryonic lethality and meiotic chromosome non-disjunction respectively, when separase function is compromised. Thus, while caspases and separases have evolved different substrate specificities associated with their specialized functions in apoptosis and cell division respectively, they appear to have retained the residual ability to participate in both processes, supporting the view that co-option of components in cell division may have led to the innovation of programmed cell suicide early in metazoan evolution.

## Introduction

Accurate segregation of chromosomes is essential for faithful transmission of the genome during somatic and germline mitotic proliferation and meiotic divisions associated with gametogenesis. Aneuploidy resulting from defective chromosome segregation can lead to a wide variety of genetic syndromes or embryonic lethality and is also associated with most malignant cells types, in some case conferring a growth advantage during cancer progression^1^. Prior to their segregation, chromosomes become aligned on the metaphase plate, where sister chromatids are held together until the onset of anaphase by the action of a ring-like structure, the cohesin complex, consisting of Scc1, Scc3, Smc1, and Smc3^2^. Sister chromatid separation at the metaphase-to-anaphase transition is initiated when Scc1 (also called Mcd1 or Rad21) is cleaved by the enzyme separase^3,4^. Separase is activated at the metaphase-to-anaphase transition as a result of degradation of its inhibitors, securin and cyclin B, by a ubiquitin protein ligase, the “anaphase promoting complex” (APC). In *C. elegans*, a single separase, encoded by the *sep-1* gene, functions during both meiosis and mitosis to promote sister chromatid separation^5^. In addition to its role in chromosome segregation, studies in *C. elegans*, Drosophila, and mammalian cells have revealed a role for separases in membrane trafficking^6–10^ Loss of separase activity in early *C. elegans* embryos also results in embryonic lethality owing to osmotic sensitivity that arises from defects in cortical granule exocytosis, as well as failure of cytokinesis, two processes that are separable from chromosome segregation^9,10^.

Separase is a member of the CD clan of cysteine proteases^4,11^. The proteases within this clan share conserved tertiary structures, arrangement of catalytic residues, and conserved motifs surrounding the catalytic residues, and apparently arose from a single evolutionary origin^12^ (Fig. 1A) The CD clan includes six distinct cysteine proteases, each that carries out unique cellular functions. These include the caspases, which are critical executors of apoptosis, or programmed cell death (PCD). When activated in cells destined to die, caspases cleave various cellular substrates, leading to the orderly dismantling of a dying cell. CED-3 is the predominant caspase in *C. elegans* responsible for nearly all of the 131 somatic and most of germline PCD during development. Three additional caspase-encoding genes encode six isoforms^13,14^, only one of which, CSP-1B, has been shown to possess proteolytic activity^15^. CSP-2 and CSP-3 instead act as negative regulators of CED-3 in the germline^16^ and soma^17^ respectively. In cells undergoing apoptosis, CED-3 is activated by trans-autoproteolysis through induced proximity mediated by the Apaf-1 homolog, CED-4^18,19^.

**Figure 1.**
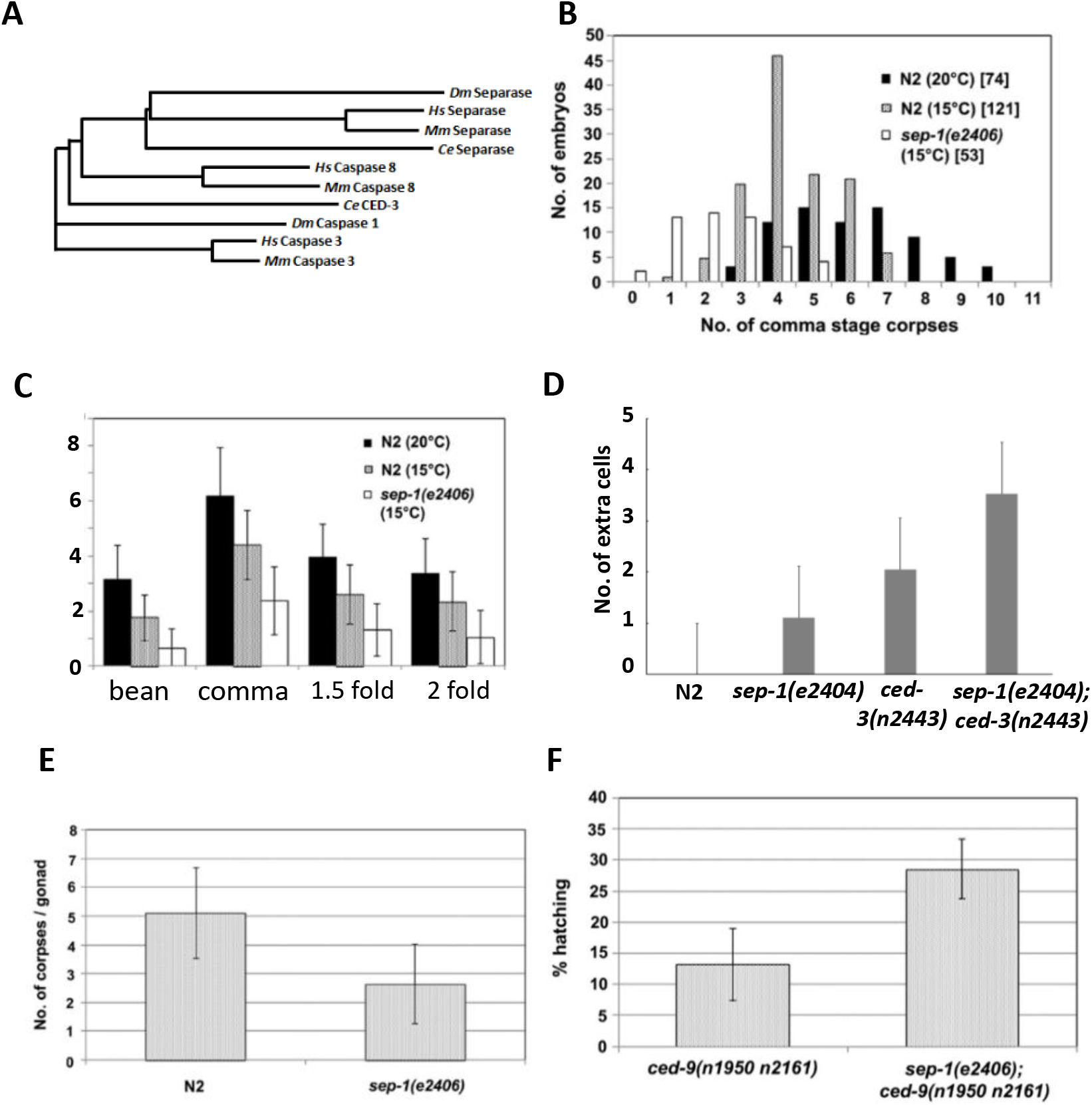
Pro-apoptotic action of separase in embryos and the germline. (A) Phylogenetic tree showing relationships of metazoan caspases and separases. *Dm*, *Drosophila melanogaster*; *Hs*, *Homo sapiens*; *Mm*, *Mus musculus*. *Ce*, *C. elegans*. (B) and (C) Subnormal numbers of cell corpses in *sep-1(ts)* embryos. In (B), cell corpses were scored at the “comma” stage at the indicated temperature. Total number of embryos scored is shown in brackets. (C) average number of cell corpses in N2 and *sep-1(e2406)* at the indicated stage and temperature. Error bars are ±SD. At least 16 embryos were scored for each stage. (D) Extra surviving cells in the anterior pharynx of *sep-1(e2406ts)* and *ced-3(n2443),* a weak allele of *ced-3,* L1 larvae of raised at the permissive temperature (15°C). (E) Decreased germ cell apoptosis in *sep-1(e2406)* mutants. Germ cell corpses were scored in both gonad arms of young adult N2 and *sep-1(e2406)* animals (50 worms were scored for each genotype) at 15°C. (F) *sep-1(e2406)* suppresses *ced-9(If)* lethality. The hatching rate of embryos produced by *sep-1(e2406); ced-9(n1950 n2161)* animals and *ced-9(n1950 n2161)* animals are shown. Total number of embryos scored is shown above each bar. Values are expressed as mean ±SD.

Mounting evidence points to regulated caspase activation in mediating other process in addition to apoptosis, including cell proliferation and differentiation ^20–24^. Some caspase targets are cell cycle regulators, including the negative regulators wee1 kinase, p27^kip1^, p21^Waf1^, the tumor suppressor Rb, and c-Abl, a kinase involved in cell cycle arrest ^25^, and there is increasing evidence that proliferation and apoptosis are coupled ^26,27^. Caspase activity is also required for T and B cell proliferation ^28,29^. The apoptotic machinery in Drosophila is also involved in spermatogenesis and oogenesis ^30–33^, and the Drosophila apical caspase *dronc* performs functions in cell migration ^34^ and proliferation ^35^. In *C. elegans*, the CED-3 caspase has been shown to cleave Dicer-1, a type III ribonuclease involved gene silencing processes by cleaving dsRNA ^36^. CED-3 was also shown to function partially redundantly with the miRNA machinery to regulate gene expression in the course of normal development by cleaving, among other proteins, LIN-28, a key regulator of developmental timing ^37,38^.

Here we present evidence that two CD clan proteases in *C. elegans*, the SEP-1 separase and the CED-3 caspase, perform shared roles in cell division and PCD. We found that *sep-1* mutants contain subnormal numbers of cell corpses, weakly enhance the PCD defect of weak *ced-3* mutants, and partially rescue the embryonic lethality of mutants lacking the PCD inhibitor CED-9/BCL-2, suggesting an involvement in PCD. We further report that CED-3 is required for the normal rate of embryonic development as well as germ cell proliferation in the hermaphrodite germline. In addition, loss of *ced-3* function enhances the cell division defect of a separase mutant in the germline and leads to meiotic chromosome non-disjunction. Our results reveal unexpected promiscuity of clan CD cysteine proteases in the processes of cell division and PCD and raise the possibility that the machinery for PCD may have arisen from components originally required for cell division.

## Results and Discussion

### SEP-1 separase is required for developmental PCD in *C. elegans*

Caspases, best known for their roles in apoptosis, and separase, an essential mitotic regulator, are members of a clade of cysteine proteases, the CD clan (Fig. 1A), and are presumed to descend from a common ancient progenitor. Caspases and separase both cleave cohesins, albeit during the very distinct cellular events of chromosome separation and apoptosis ^4,25,39–41^. We sought to investigate whether their potential functional overlap might be more than superficial by assessing whether SEP-1 separase is discernibly required for normal PCD. Loss of SEP-1 results in early embryonic lethality prior to onset of the developmental PCD as a result of osmotic sensitivity stemming from defects in eggshell formation as well as failure of cytokinesis and proper chromosome segregation^42 5,9,10^ To assess later roles of SEP-1, we took advantage of the temperature-sensitive *sep-1(e2046)* mutation, a missense allele (C450Y) that primarily affects vesicle trafficking and cortical granule exocytosis and that has minimal effect on chromosome segregation^10^. *sep-1(e2046)* animals show fully penetrant embryonic lethality at 22°C, but are viable at 15°C. We found that the *sep-1(ts)* mutant grown at permissive temperature showed a highly significant reduction in the number of apoptotic corpses (average of 2.4 ± 1.2 corpses; n = 53; p = 6.4 × 10^−15^) at the “comma” stage of embryogenesis compared to that in wild type (N2) embryos (average of 4.4 ± 1.2; n = 121; Fig. 1B). This decrease in the number of cell corpses does not appear to result from a delay in the execution of embryonic PCD, as we observed significantly fewer corpses throughout all late stages of *sep-1(ts)* embryos (Fig. 1C).

If the reduction in cell corpses reflects a *bona fide* block to PCD in some cells, it would be expected to result in supernumerary surviving cells, as is conveniently assayed by quantifying nuclei in the anterior pharynx ^43^. Consistent with the decrease in cell corpses, we found that *sep-1(ts)* mutants grown at 21°C contained extra surviving nuclei in the anterior pharynx of L1 larvae (average of 1.1 ± 0.66 extra cells; n=19; p = 8.42 × 10^−7^) compared to the laboratory reference strain N2 (Fig. 1D). This value is similar to that seen in some weak *ced-3(−)* mutants ^44^. In addition, we found that *sep-1(e2046)* grown at 21°C significantly enhances the cell death defect of *ced-3(n2443)*, a weak reduction-of-function allele, resulting in an average of 3.53 ± 0.91 extra nuclei in the anterior pharynx (n=19; p = 2.86 × 10^−6^), compared to an average of 2.05 ± 0.71 in the *ced-3(n2443)* single mutant. Further supporting a role for SEP-1 in PCD, we found that the number of cell corpses in the adult germline is reduced approximately two-fold (2.6 ± 1.4 corpses; p = 6 × 10^−11^) in the *sep-1(ts)* mutant grown at permissive temperature compared to that in wild-type (Fig. 1E).

The observation that a *sep-1* mutation even at permissive temperature appears to suppress the death of individual cells that are normally developmentally programmed to die raises the question of whether SEP-1 might be broadly involved in regulating PCD. The loss of CED-9/Bcl-2, a major apoptotic suppressor, results in maternal-effect lethality caused by massive ectopic PCD in embryos ^45^. We found that the *sep-1(e2046)* mutation suppresses the maternal-effect lethality of *ced-9(n1950 n2161*) loss-of-function mutant: while 13.2 ± 5.8% (n = 3,036 total embryos from 39 worms) of *ced-9(n1950 n2161*) loss-of-function mutant embryos cultured at 15°C hatch, a significantly higher number of *sep-1(e2046)*; *ced-9(n1950 n2161*) double mutant survive to hatching (28.6 ± 4.8%; n = 5,626 total embryos from 40 worms; p=1.5 × 10^−13^)(Fig. 1F). Further, while the small number of hatching *ced-9(−)* mutants never developed past the early L1 stage, we found that approximately half of the arrested *sep-1(−); ced-9(If)* double mutant larvae grew to a larger size and appeared to have progressed further (late L1 or L2) before arresting.

While CED-3 is the predominant caspase in mediating apoptosis, these findings implicate a requirement for SEP-1 in PCD. We note that these experiments were performed under conditions in which SEP-1 is virtually fully functional for its known separase activities, at least with respect to its role in cortical granule exocytosis, cytokinesis, and chromosome segregation. The early embryonic lethality in strongly affected *sep-1(−)* mutants precluded us from scoring PCD and extra cells in animals at the non-permissive temperature for *sep-1(ts).* Although the effects we have observed for the *sep-1(−)* are relatively mild, is conceivable that separase could actually play a major essential role in the execution of PCD that was heretofore masked owing to its essential requirement in mitosis and early embryonic viability.

### CED-3, but not PCD, is required for the normal rate of embryonic development

Given that separase and apoptotic caspases belong to same clade, the CD clan cysteine proteases, coupled with the evidence presented above that separase may possess pro-apoptotic activity, it was of interest to ask whether the predominant pro-apoptotic caspase in *C. elegans*, CED-3, might function in essential non-apoptotic roles performed by separase. The loss of *ced-3* caspase function eliminates nearly all PCD in *C. elegans*^14^; however, it does not lead to any debilitating or other overt phenotypes. This finding originally prompted the view that CED-3 caspase functions primarily or exclusively in PCD. While CED-3 is not essential for viability or fertility in *C. elegans*, we found that the *ced-3(n717)* mutation results in substantially slower embryonic development (Figure 2A), and hence an increased time to hatching by ~20% or ~120 min, compared to wildtype N2 (p = 0.003), suggestive of a broader role for CED-3. To rule out specific effects of the *ced-3(n717)* mutant background on developmental rate, we analyzed six additional alleles of *ced-3*, of which three (*n718*, *n2454* and *n2442)* show a strong defect in PCD, and the remainder (*n1040*, *n2877* and *n2921)*, a more moderate defect in PCD. Irrespective of their effect on PCD, all six alleles cause a delay in embryonic development similar to that seen in *ced-3(n717)* mutants (Figure 2A). This slower developmental rate is not attributable to the absence of PCD, as a mutation in *ced-4*, the gene encoding *C. elegans* pro-apoptotic Apaf1 ^46^, which also abolishes PCD, had no discernible effect on the rate of embryogenesis. These observations suggest that *ced-3* performs functions that are important in normal progression through embryogenesis distinct from its function in activation of PCD.

**Figure 2.**
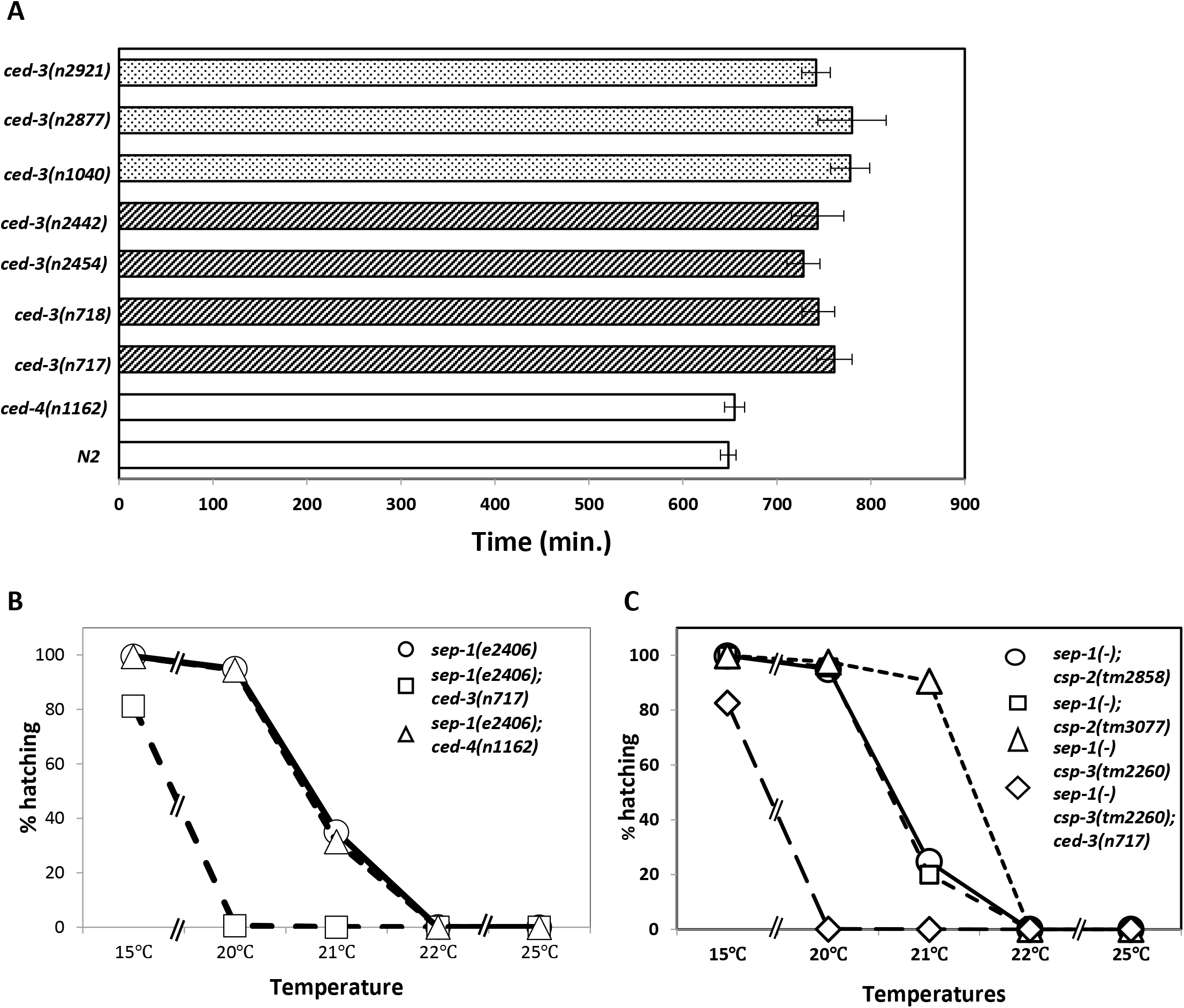
Delayed embryonic development in *ced-3* mutants. (A) Slowed embryonic development in *ced-3* mutants. Embryos of the indicated genotype were allowed to develop at 22.5±0.5°C and the time from 2-cell to hatching was measured. Stippled bars: *ced-3* alleles that show a moderate defect in PCD (33). Striped bars: *ced-3* alleles that show a strong defect in PCD. (B) Embryonic survival. Embryos of the indicated genotype and temperature were scored for hatching as the measure of embryonic viability. Each data point was obtained from >154 embryos. (C) Partial suppression of *sep-1(−)* embryonic lethality by *csp-3(−)*. Embryos of the indicated genotype were scored as in A. Each data point was obtained from >1000 embryos.

### Vital embryonic role of CED-3 is revealed when SEP-1 function is reduced

Mutations in *sep-1* affect egg shell formation, cause defects in cytokinesis and chromosome separation, and block mitosis, in all cases leading to embryonic lethality ^5,9,10,47^. We found that *ced-3(−)* mutations greatly enhance the embryonic lethality of *sep-1(ts)* mutants grown at a fully permissive or semi-permissive temperature range of 15-22°C (Fig. 2B). This effect was most striking at an intermediate temperature of 20°C: while *sep-1(ts)* single-mutant embryos nearly always (~95%; n = 2,564) survive, those also lacking CED-3 function invariably die (<1% viable; n = 733). Given that *sep-1(e2406)* is a phenotypically complicated allele, we also compromised SEP-1 function by RNAi. RNAi of *sep-1* in wild type (N2) background results in high lethality. While the embryonic lethality progressively decreases with increasing dilution in N2 background it remains higher in the *ced-3* mutant background (Figure S1A). In addition to CED-3, the CSP-1B caspase, one of three isoforms encoded by *csp-1*, has been implicated in regulating a subset of embryonic cells destined to die ^48^. We found that knockdown of *csp-1* by RNAi had no effect on viability of *sep-1(ts)* mutant embryos (not shown); hence this effect is not general to caspases. More importantly, the greatly enhanced lethality is not related to abrogation of PCD *per se*, as a *ced-4(−)* mutation that eliminates virtually all PCD showed no enhancement of lethality in the *sep-1(ts)* mutant (Fig. 2B).

Two non-catalytic caspase homologs, CSP-2 and CSP-3, function to buffer caspase activity in living cells by inhibiting CED-3 in the germline and soma respectively. We found that removal of CSP-3, the soma-specific caspase inhibitor ^17^, suppresses the high embryonic lethality (65%) seen with the *sep-1(ts)* mutant grown at a semi-permissive temperature of 21°C to 9% lethality (p < 10^−4^) (Figure 2C). This strong suppression of lethality requires CED-3 function: the *sep-1(ts); csp-3(−); ced-3(−)* triple mutant shows 100% lethality at 20°C, which is nearly fully permissive for the *sep-1(ts)* mutation alone (Fig. 2C). As expected, loss of the germline-specific CED-3 inhibitor CSP-2 ^16^ does not suppress this lethality (Figure 2C).

### Synergistic requirement for CED-3 and SEP-1 in osmotic integrity and chromosome segregation

The apparently shared requirement for CED-3 and SEP-1 in PCD and viability raises the possibility that CED-3 might also affect chromosome segregation when separase activity is attenuated. To follow chromosome separation in living embryos, we observed fluorescently tagged chromosomes during cleavage of the zygote (Fig. 3). The *sep-1(ts)* strain shows defects in vesicle trafficking and exocytosis that result in cytokinetic failure, impaired egg shell resulting in osmotic sensitivity, and chromosome segregation and centriole disengagement, defects which are separable from each other ^9,10,47,49^. To ameliorate effects on osmotic sensitivity we analyzed embryos in osmotically balanced egg salt buffer. We found that nearly all mutant embryos carrying a temperature-sensitive mutation in *sep-1* progress beyond the first division when incubated at a semi-permissive temperature of 21°C. Under these conditions, the average time required for the transition from maximum alignment of metaphase chromosomes to initiation of chromosome segregation in wild-type was ~101 ± 14 sec (n = 20; Fig. 3A and B). While chromosome separation also succeeded in nearly all (95%; n = 21) *sep-1(e2406ts)* mutants at this temperature, the average time from maximum metaphase alignment to initiation of separation increased significantly, by nearly 2-fold, to ~3 min (p = 3.2 × 10^−7^) (Fig. 3A and B). As expected, chromosome separation and cytokinesis were always successful in *ced-3(−)* single mutants and the time for this event was virtually indistinguishable from that in wild-type (105 ± 18 sec; n = 20; Fig. 3B). In contrast, we found that nearly half (43%; n = 31) of *sep-1(ts)*; *ced-3(−)* double mutant zygotes completely failed to undergo chromosome separation or cytokinesis (Fig. 3A). In the profoundly defective embryos, no prominent pseudocleavage furrow was observed and the embryo filled the eggshell. This suggests that the osmotic integrity of the eggshell in the double mutant was compromised under the same conditions in which the *sep-1(e2406ts)* single mutant embryos were able to complete zygotic cell division. We also found that chromatin condensation was affected, multipolar chromatin bridges were observed, and chromosomes remained in the center of the cell indefinitely (>40 min). This profound enhancement of the chromosome segregation defects is not attributable to the block in germline PCD in the *ced-3(−)* mothers of the embryos *per se*, as mutations in *ced-4*, which also block virtually all PCD, do not enhance, and in some cases actually suppress, chromosome segregation phenotypes associated with loss of *sep-1* function (see below). These findings implicate a partially redundant role for the pro-apoptotic caspase CED-3 in conjunction with separase during eggshell formation, cytokinesis, and chromosome segregation.

**Figure 3.**
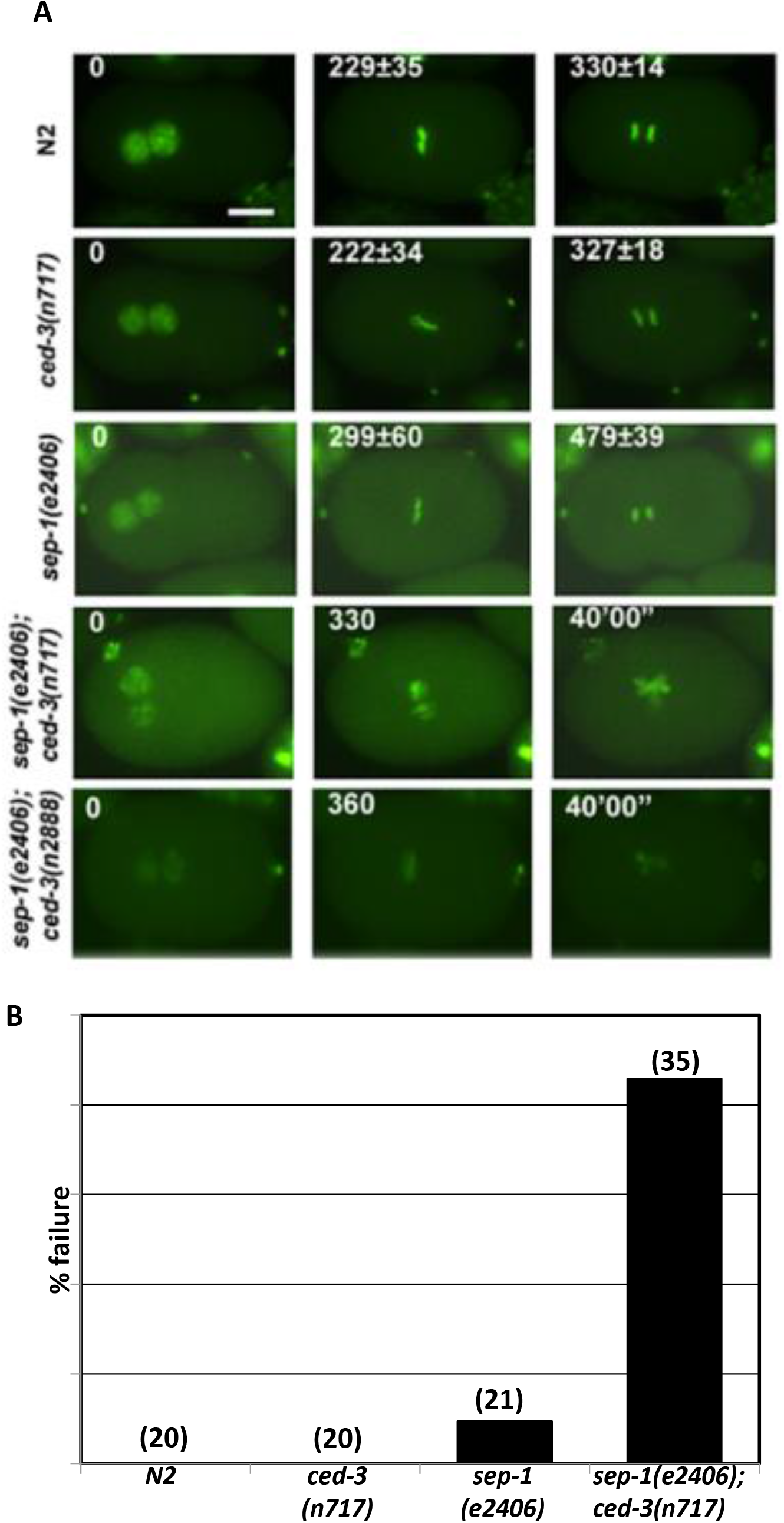
CED-3 enhances early cell division defects in separase mutants. (A) Progression of GFP-labeled mitotic chromosomes during zygotic cleavage. One-cell embryos of the indicated genotype, carrying the HIS-2B::GFP marker, are shown at pronuclear meeting (defined as t=0; left panels), maximum metaphase alignment (middle panels), and immediately following separation of chromosomes (right panels, with the exception of the *sep-1(e2406); ced-3(n717)* embryo, which failed in chromosome separation). Embryos were dissected from young adults that had been pre-incubated at 21°C for 90 mins and allowed to undergo cleavage at the same temperature. The time following pronuclear meeting is indicated in seconds, with the standard deviation reported for all embryos (see B) that underwent chromosome segregation with the exception of the last *sep-1(−)*; *ced-3(−)* panel, in which time in indicated in minutes. Scale bar, 10 μm. The arrow and arrowhead indicate pseudocleavage and cleavage furrows respectively which are absent in the double mutant embryos. The asterisks show the chromatin condensation and separation defects in the last two panels of the double mutant embryos. (B) Average time for chromosome segregation in all embryos of the indicated genotype (n = 20 per genotype) in which separation of chromosomes was successful. The time from metaphase alignment to the start of chromosome segregation, with standard deviation, is indicated. For all experiments, embryos were imaged every 30 sec with a 500 ms exposure time.

**Figure 4.**
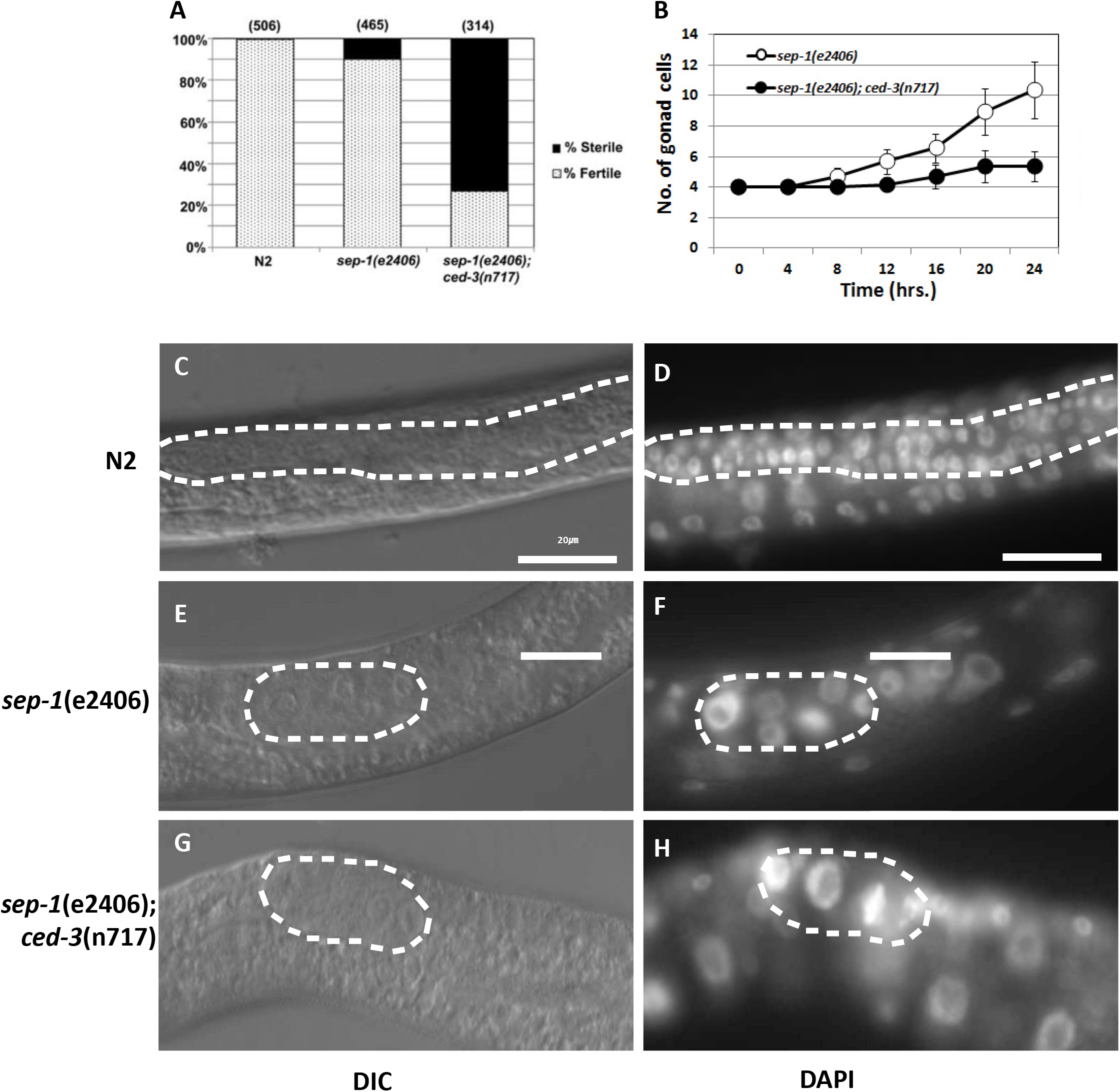
Synergy of CED-3 and SEP-1 in gonadal cell proliferation. (A) % sterility of adults grown at 15°C. (B) Quantitation of gonadal (both somatic and germline) nuclei in larvae of the indicated genotype. Synchronized L1 worms (n = 50 for each data point) were grown at 25°C, and samples collected every 4 hrs. Larvae were fixed in Carnoy’s solution, stained with DAPI, and gonadal cells counted. (C), (D) and (F), DIC images of N2, *sep-1(e2406)* and *sep-1(e2406); ced-3(n717)*, respectively, 24 hours after initiation of feeding of synchronized L1 larvae grown at 25°C. (D), (F) and (G), corresponding DAPI images of the same animals.

### Requirement for CED-3 activity in germ cell proliferation

The mechanisms that regulate proliferation in the *C. elegans* germline are distinct from those acting in the embryo ^50,51^. We therefore asked whether the requirement for CED-3 extends to germ cell proliferation. *sep-1(e2406ts)* mutants show a low level of sterility at a permissive temperature of 15°C (9.9%; n=465). We found that this sterility is dramatically enhanced (to 72.9%; n = 314) by simultaneous removal of *ced-3* activity (Figure 3A). Examination of the sterile adult animals showed that they contained underproliferated germlines and frequent abnormal oocytes (not shown). Analysis of post-embryonic germline development in synchronized worms cultured at a non-permissive temperature of 25°C starting immediately after hatching revealed that the block in germ cell proliferation resulting from loss of SEP-1 is greatly exacerbated by removal of CED-3 (Figure 3). At hatching, the gonad primordium of wild-type worms contains 4 nuclei, which increases to an average of 44.2 ± 3.6 (n = 30) nuclei per gonad after 24 hours, (Suppl. Figure S1). Under these conditions, *sep-1(ts)* animals contained an average of 10.4 nuclei per gonad (n = 50 animals), all of which were somewhat larger and stained more intensely with DAPI, revealing that removal of SEP-1 function strongly blocks cell proliferation in the gonad (Figure 3B, S1-D). We found that elimination of CED-3 function in the *sep-1(−)* mutant virtually completely blocks all proliferation of the initial 4 gonadal cells (the two somatic gonad precursor cells as well as the germline precursor cells) at hatching, with an average of only 5.4 nuclei/gonad (n=50) (Figure 3B,S1-F). The very strong DAPI staining and increased size of the chromosome aggregates seen in *sep-1(ts); ced-3(−)* double mutants suggest that all gonadal nuclei have apparently continued to undergo multiple rounds of DNA replication without karyokinesis (Figure S1).

### CED-3 is required for normal X-chromosome segregation in the germline

In *C. elegans*, sex is determined by the number of X-chromosomes relative to the number of autosomes, such that XX embryos develop into hermaphrodites and XO embryos become males ^52^. A number of mutants that show X-chromosome nondisjunction result in a <u>h</u>igh <u>i</u>ncidence of <u>m</u>ales (Him) phenotype ^53^. We found that *sep-1(ts)* single mutant animals grown at the permissive temperature exhibit a nearly nine-fold increased rate of male production (1.1%; p < 10^−4^) compared to wild-type, consistent with the role of SEP-1 in chromosome segregation. Similarly, we found that two different alleles of *ced-3* result in a statistically significant (p <0.01) increase in male production at 15°C by as much as four-fold (0.48% males) compared to the wildtype (0.12%) (Table 1), implicating the CED-3 caspase in proper chromosome disjunction during meiosis. Further, we found that eliminating both *ced-3* and *sep-1* function together results in a strongly synergistic effect: *sep-1(ts); ced-3(−)* animals show a >50-fold enhancement (6.2%; p < 10^−4^) in the rate of male production over wild-type. These results suggest that CED-3 and SEP-1 might cooperate in meiotic chromosome segregation.

**Table 1.**
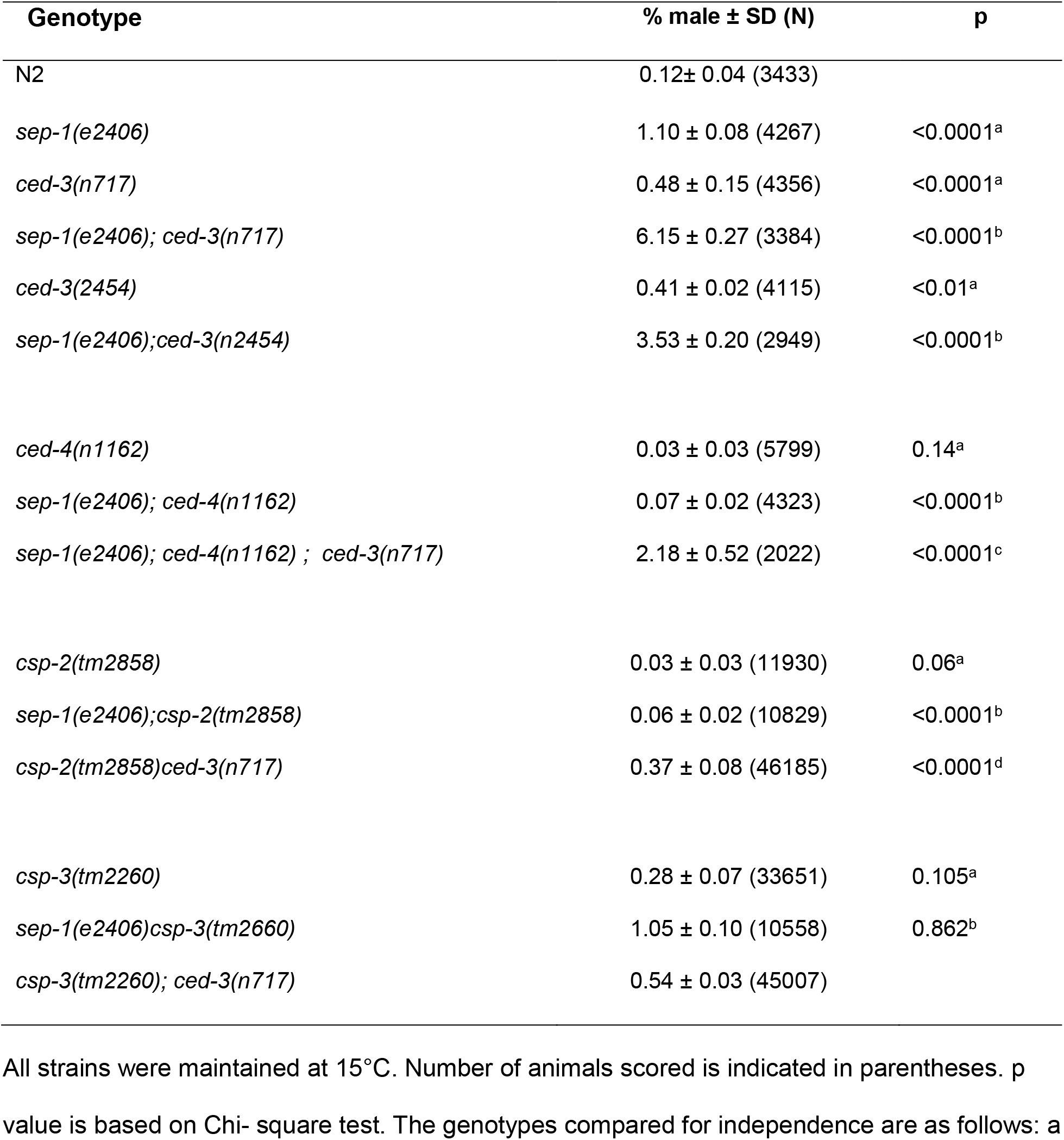
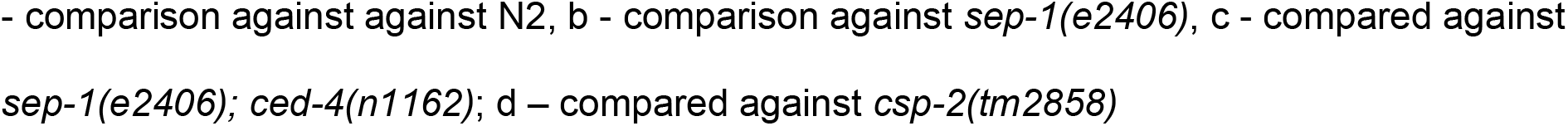
Quantitation of X-chromosome non-disjunction.

| Genotype | % male ± SD (N) | p |
| --- | --- | --- |
| N2 | 0.12± 0.04 (3433) | |
| *sep-1(e2406)* | 1.10 ± 0.08 (4267) | <0.0001<sup>a</sup> |
| *ced-3(n717)* | 0.48 ± 0.15 (4356) | <0.0001<sup>a</sup> |
| *sep-1(e2406); ced-3(n717)* | 6.15 ± 0.27 (3384) | <0.0001<sup>b</sup> |
| *ced-3(2454)* | 0.41 ± 0.02 (4115) | <0.01<sup>a</sup> |
| *sep-1(e2406);ced-3(n2454)* | 3.53 ± 0.20 (2949) | <0.0001<sup>b</sup> |
| *ced-4(n1162)* | 0.03 ± 0.03 (5799) | 0.14<sup>a</sup> |
| *sep-1(e2406); ced-4(n1162)* | 0.07 ± 0.02 (4323) | <0.0001<sup>b</sup> |
| *sep-1(e2406); ced-4(n1162) ; ced-3(n717)* | 2.18 ± 0.52 (2022) | <0.0001<sup>c</sup> |
| *csp-2(tm2858)* | 0.03 ± 0.03 (11930) | 0.06<sup>a</sup> |
| *sep-1(e2406);csp-2(tm2858)* | 0.06 ± 0.02 (10829) | <0.0001<sup>b</sup> |
| *csp-2(tm2858)ced-3(n717)* | 0.37 ± 0.08 (46185) | <0.0001<sup>d</sup> |
| *csp-3(tm2260)* | 0.28 ± 0.07 (33651) | 0.105<sup>a</sup> |
| *sep-1(e2406)csp-3(tm2660)* | 1.05 ± 0.10 (10558) | 0.862<sup>b</sup> |
| *csp-3(tm2260); ced-3(n717)* | 0.54 ± 0.03 (45007) | |

All strains were maintained at 15°C. Number of animals scored is indicated in parentheses. p value is based on Chi- square test. The genotypes compared for independence are as follows: a
- comparison against against N2, b - comparison against *sep-1(e2406)*, c - compared against *sep-1(e2406); ced-4(n1162)*; d – compared against *csp-2(tm2858)*

Just as CSP-3 functions to buffer CED-3 pro-apoptotic activity in the soma, CSP-2 performs an analogous role in the germline. We found that removal of CSP-2, but not CSP-3, suppresses the Him phenotype of *sep-1(−)* mutants (0.06% *versus* 1.1% male production; p <10^− 4^; Table 1), consistent with an increase in CED-3 activities that might compensate for the lack of SEP-1. In contrast, the increased production of males in *ced-3(−)* single mutants is not significantly suppressed by elimination of CSP-2, as seen in the *csp-2(−)*; *ced-3(−)* double mutant (Table 1), implying that CSP-2 functions through CED-3 in this process. Further, elimination of CSP-2 appears to decrease the naturally observed Him phenotype compared to that of wild-type (0.03% *vs.* 0.12%; p = 0.06), suggesting that elevated CED-3 levels resulting from the absence of CSP-2 may be sufficient to diminish the natural X non-disjunction seen in wild-type animals containing normal CED-3 and SEP-1.

In sharp contrast, a *ced-4(−)* mutation, which blocks PCD as effectively as a strong *ced-3(−)* mutation, has no effect on, or perhaps even slightly reduces chromosome non-disjunction compared to wild-type (0.03%; p = 0.14). Further, the increased X chromosome non-disjunction seen in the *sep-1(ts)* mutant is suppressed by the *ced-4* mutation (0.07%; p < 10^−4^ for *sep-1(ts) vs. sep-1(ts); ced-4(−)*); i.e., removal of CED-4 counteracts the effect of debilitating separase function. CED-4/Apaf1 functions in apoptosis by binding to and directing CED-3 to the apoptotic pathway, a process that is very active in the germline ^54^. We posit that removal of CED-4 may result in elevated levels of uncomplexed CED-3 that becomes available to participate in chromosome separation. Supporting this notion, we found that the suppression of meiotic non-disjunction by *ced-4(−)* is eliminated by removal of *ced-3(+)* activity (compare *sep-1(e2406); ced-4(n1162)* to *sep-1(e2406); ced-4(n1162); ced-3(n717)*; p < 10^−4^; Table 1). Collectively, these results implicate the CED-3 caspase in meiotic chromosome segregation and suggest that CSP-2 and CED-4 antagonize its action in this process.

## Discussion

This study and other recent reports reveal multiple functions of proteins that regulate cell death and also perform other vital functions in surviving cells ^21,55–57^. Pro-apoptotic CED-4/Apaf1 has been reported to mediate DNA-damage-induced cell-cycle arrest at S phase ^55^. CED-4 has also been shown to be required for regulation of cell size ^21^. Caspases have been implicated in multiple cellular processes not related to cell death, including terminal differentiation, activation, proliferation, and cytoprotection ^56^. Caspase-2 was also found to be involved in maintenance of the G2/M DNA damage checkpoint and DNA repair ^57^. We note that caspase-2 is the only caspase that is constitutively present in the cell nucleus and is closely related to the *C. elegans* CED-3 ^44^. In the worm, a consequence of abrogation of programmed germ cell death by *ced-3* has been a reduction in quality of oocytes in aging worms due to a reduction in allocation of resources to developing oocytes ^58^. CED-3 proteolytic activity, in conjunction with the miRNA silencing machinery^38^ and the N-end rule degradation machinery^37^, plays a role in regulating the robust expression of many genes during development. Dicer, a type-III ribonuclease is converted into a pro-apoptotic DNAase by CED-3^36^ and consequentially might affect gene silencing by non-coding RNAs.

Like caspases, separase is predicted to have multiple cleavage substrates based on a consensus cleavage sequence ^40,59^. Separase has been reported to cleave a kinetochore/spindle protein called Slk19 ^60^. Some of the potential separase substrates do not have mitotic functions and may provide clue to its role in other processes. Multiple studies have raised the possibility of a role for separase in apoptosis-related functions. Separase has been reported to cleave the yeast Scc1 to amplify of apoptotic signal initiated by H_2_O_2_ ^61^. In another study, separase inhibition was found to be required for the survival of murine embryonic cells ^62,63^. A constitutively active separase mutant leads to apoptotic cell death during embryonic development ^62,63^. These results, coupled with our findings, suggest that separase may play a caspase-like executioner role and cleave multiple protein substrates during apoptosis.

The interplay between the cell death machinery and essential cellular activities is consistent with the notion that some of the molecular components regulating normal cellular processes may have been recruited into the complex apparatus that orchestrates the demise of cells ^64^. The action of separase in mitosis is likely to be among its most ancient functions, as essentially all eukaryotes require this enzyme to direct chromosome segregation at mitosis ^2,5^. Apoptosis was a major innovation associated with the emergence of multicellularity in metazoans ^64,65^. The essential role of a cysteine protease that is required for cell division in developmentally programmed cell death supports the view that the apoptotic machinery arose as a consequence of co-option of basic cellular machinery involved in mitosis. Such co-option of essential cellular proteases as cellular executioners would presumably have been accompanied by the coevolution of buffering mechanisms (as provided, for example, by the CSP-2 and −3 inhibitors) that allow fine-tuning of enzymatic activity required to mediate essential cellular functions. Discovering the contribution of essential genes to PCD is complicated by the fact that their depletion often leads to lethality. The Janus-like behavior of SEP-1 might be representative of many other unexplored apoptotic functions of the essential components in the cellular toolkit.

The tight regulation of cell proliferation and cell survival is of fundamental importance to normal development and homeostasis. The notion of a connection between cell death and proliferation derives in part from evidence that apoptosis is a consequence of dysregulation of the cell-cycle machinery ^26^. Disrupting or bypassing cell-cycle checkpoints frequently leads to increased apoptotic potential instead of increased proliferation. The involvement of common effector molecules CED-3 and SEP-1 at the interface between mitosis and apoptosis points to a further link between these two key cellular processes and provides a framework for understanding the complex signaling pathways that coordinate cell proliferation and cell death. These findings also raise a note of caution with the design of anti-tumor drug therapies targeted to separase activity; as we have observed, inhibition of this enzyme with the goal of blocking its mitosis-directing function might simultaneously prevent the major anti-tumor defensive mechanism, apoptosis, thereby potentially confounding such a strategy.

## Materials and methods

### Strains and culturing

*C. elegans* was cultured at 20°C by standard procedures ^66^, unless otherwise noted. Temperature-sensitive strains were maintained at 15°C. Strain N2 Bristol variety was used as the wild-type. *sep-1*(*e2406*) was isolated from strain WH216 *sep-1(e2406)*/*hT2[bli-4(e937)let-?(q782) qIs48]* (I;III), obtained from the *Caenorhabditis* Genetics Center (CGC), by backcrossing five times to generate JR3391. AZ212 *unc-119(ed3); ruls32[unc-119(+) pie-1::GFP::H2B]* was obtained from the CGC and used to construct JR3304 *ruIs32; ced-3(n717),* JR3388 *sep-1*(*e2406*)*; ruIs32*, JR3389 *sep-1(e2406*)*; ruIs32; ced-3* (*n717*) by standard methods. *csp-2(tm2858), csp-2(tm3077), csp-3(tm2260)*, and *csp-3(tm2486);csp-2(tm3077)* were gifts from Dr. Ding Xue ^16,17^. The various *ced-3* alleles were a gift from Dr. H. Robert Horvitz ^44^.

### Time-lapse analysis of chromosome segregation

*sep-1(e2406)* and *sep-1(e2406); ced-3(n717)* embryos expressing GFP-tagged histone 2B (*ruls32*) were obtained from adult worms that had been pre-incubated at 21°C for 90 min. Early one-cell embryos collected from dissected worms were mounted on agarose pads in egg salts buffer. Fluorescence time-lapse images were acquired at 24±1°C using a Nikon Eclipse Ti microscope controlled by NIS Elements AR software. Specimens were illuminated with an X-cite light source attenuated to 20% using a GFP filter (480/40 bandpass excitation filter). All images were obtained with a Hamamatsu CMOS sensor using a 20X objective. Images were acquired every 30 sec. by scanning through five to eight focal planes per time point.

## Acknowledgments

We thank S. Shaham for the *csp-2* and *-3* mutants, H. R. Horvitz for the *ced-3* alleles, and S. Martin for the pGEX-CED-3 expression vector. Some nematode strains used in this work were provided by the Caenorhabditis Genetics Center, which is funded by the NIH National Center for Research Resources (NCRR). This work was supported by grants from the NIH (#HD081266 and #HD082347) to J.H.R.

## Author contributions

PYJ was involved in the design of all experiments and obtained and analyzed the data presented in the manuscript. AK performed initial experiments implicating a role for SEP-1 in PCD. PJ was involved in experimental design, data analysis, and writing of the manuscript. JHR directed the project and was involved in the concept, experimental design, and data analysis of the project and writing of the manuscript.

## Competing interests

The authors declare no competing interests

---

**Supplemental Figure S1.**
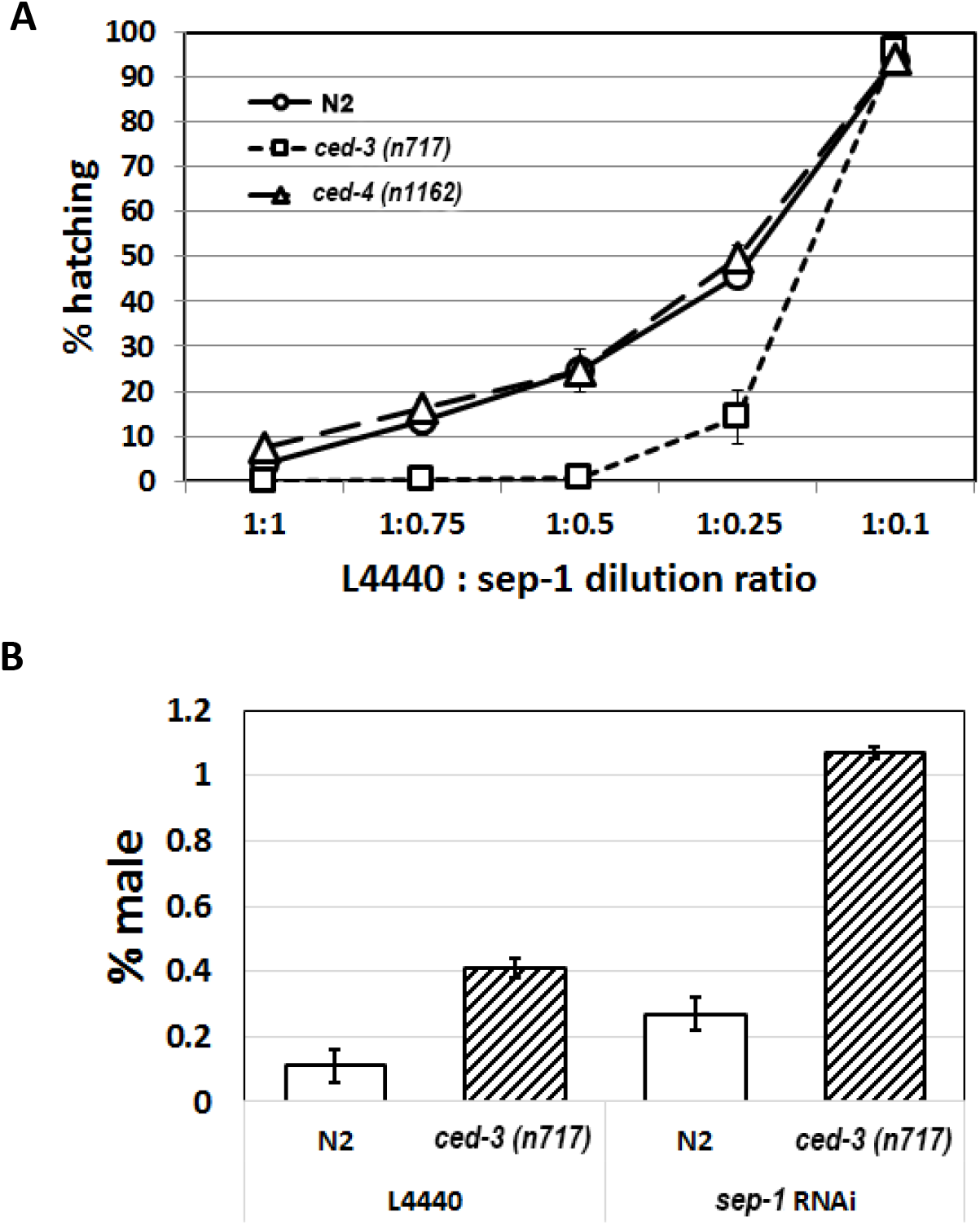
*ced-3* enhances reduction of function defects of *sep-1* RNAi treated animals. (A) Embryonic lethality of embryos from parents fed with HT115 E.coli strain expressing dsRNA against sep-1 diluted with bacteria carrying an empty vector. (B) % of males among the surviving progeny of L4 parents fed diluted sep-1RNAi bacteria.

